# Longitudinal epi-transcriptome profiling reveals the crucial role of m^6^A in prenatal skeletal muscle development of pigs

**DOI:** 10.1101/2019.12.27.888560

**Authors:** Xinxin Zhang, Yilong Yao, Jinghua Han, Yalan Yang, Yun Chen, Zhonglin Tang, Fei Gao

**Author notes:** Correspondence should be addressed to Z.T. and F.G. These authors contributed equally to this work (Xinxin Zhang, Yilong Yao).

## Abstract

**Background:** N6-methyladenosine (m^6^A) is the most abundant RNA modification and essentially participates in the regulation of skeletal muscle development. However, the status and function of m^6^A methylation in prenatal myogenesis remains unclear now.

**Results:** In our present study, we first demonstrate that chemical suppression of m^6^A and knockdown METTL14 significantly inhibit the differentiation and promote the proliferation of C2C12 myoblast cells. The mRNA expression of m^6^A reader protein IGF2BP1, which functions to promote the stability of target mRNA, continually decreases during the prenatal skeletal muscle development. Thereafter, profiling transcriptome-wide m^6^A for six developmental stage of prenatal skeletal muscle, which spanning two important waves of pig myogenesis, were performed using a refined MeRIP sequencing technology that is optimal for small-amount of RNA samples. Highly dynamic m^6^A methylomes across different development stages were then revealed, with majority of the affected genes enriched in pathways of skeletal muscle development. In association with the transcriptome-wide alterations, transcriptional regulatory factors (MyoD) and differentiated markers (MyHC, MYH1) of muscle development are simultaneously regulated with m^6^A and IGF2BP1. Knockdown of IGF2BP1 also suppresses myotube formation and promotes cell proliferation.

**Conclusions:** The present study clarifies the dynamics of RNA m^6^A methylation in the regulation of prenatal skeletal muscle development, providing a data baseline for future developmental as well as biomedical studies of m^6^A functions in muscle development and disease.

## Introduction

Understanding the development of skeletal muscle is crucial to unravel the molecular basis of formation and diseases of skeletal muscle [1, 2]. The formation of skeletal muscle is regulated not only by genetic factors but also by epigenetic factors [3, 4]. Among these factors, N6-methyladenosine (m^6^A) is an epigenetic factor newly discovered in recent years [5, 6]. A dynamic regulation of m^6^A during development is achieved through interplay among m^6^A methyltransferases (METTL3/METTL14/WTAP) and demethylase (FTO/ALKBH5) [7–9]. Despite that a previous study revealed the transcriptome-wide m^6^A profile of porcine muscle tissue, to date, very limited studies have been accomplished for the role of m^6^A in prenatal myogenesis [10]. Thereby, it is of great significance to understand the dynamic regulation of m^6^A throughout the myogenesis process. In addition to the difficulty of sampling prenatal human tissue, lack of proper technology that can analyze m^6^A from a small amount of RNAs also hinders m^6^A screening of prenatal myogenesis [11]. Alternatively, animal models can be used to study the underlying mechanisms. Due to the similarity with humans in anatomy, physiology/pathology and even genome [12, 13], pig is an ideal model animal for skeletal muscle research. Like human, the myogenesis process of pig takes place in two distinct waves before birth, which ultimately determines the total number of fibers (TNF). The first wave happens at 35 to 64 dpc (days post coitus) to form the primary myofibers, whereas the second wave happens at 54–90dpc to form the secondary myofibers [14]. The profiling of m^6^A maps of adult porcine muscle tissues from three different breeds [10] provided a valuable reference for further study of using pig as a model to study the dynamic regulation and biological significance of m^6^A for myogenesis.

In present study, we confirmed that the chemical inhibition as well as the knock down of methionine adenosyltransferase could lead to impair cell differentiation and promote proliferation of C2C12 myoblast cell. A refined MeRIP (m6A-specific methylated RNA immunoprecipitation with nextgeneration sequencing) method [11] was then applied to profile the m^6^A epi-transcriptomes of six prenatal life stages (33dpc, 40dpc, 50dpc, 60dpc, 70dpc, 95dpc) of pig skeletal muscle tissues, spanning two important waves of pig myogenesis. Inter-stage fold changes of m^6^A and gene expression was significantly correlation. Reader protein IGF2BP1 (insulin like growth factor 2 mRNA binding protein 1) expression continually decrease during prenatal skeletal muscle development, and siRNA-mediated knockdown of IGF2BP1 inhibited myotube formation and promoted proliferation in myoblast. We found that MyoD, MyHC and MYH1 gene expression was synergistically regulated by m^6^A and IGF2BP1. In conclusion, our result uncovered that m^6^A is a crucial epigenetic factor in porcine prenatal myogenesis.

## Results

### m^6^A regulates the proliferation and differentiation of C2C12 myoblast

To investigate the role of m^6^A modification in myogenesis, we first used Cycloleucine, a competitive and reversible inhibitor of methionine adenosyltransferase [15], to inhibite m^6^A level in C2C12 myoblast. Two doses of Cycloleucine treatments suppressed myotube formation, as seen by the morphology of Myosin immunofluorescence staining (Fig. 1a); and the expression of transcriptional regulatory factors (MyoD) and differentiated markers (MyHC) [16, 17] was significantly down-regulated at mRNA level (Fig. 1b, c), indicating the Cycloleucine can inhibited the C2C12 myoblast differentiation. To observe the function of m^6^A in myoblast proliferation, we used 5-ethynyl-2-deoxyuridine (EdU) staining to analyze the C2C12 cell proliferation. The EdU staining assay showed that two doses of Cycloleucine treatments significantly increased EdU incorporation (Fig. 1d, e), suggesting the involvement of m^6^A in regulating C2C12 myoblast proliferation. Moreover, knockdown of methyltransferases METTL14 resulted in the same phenotype as Cycloleucine treatments (Fig.1f-j). Taken together, these observations suggest that m^6^A played an important role in myogenesis by regulating myoblast proliferation and differentiation.

**Fig. 1.**
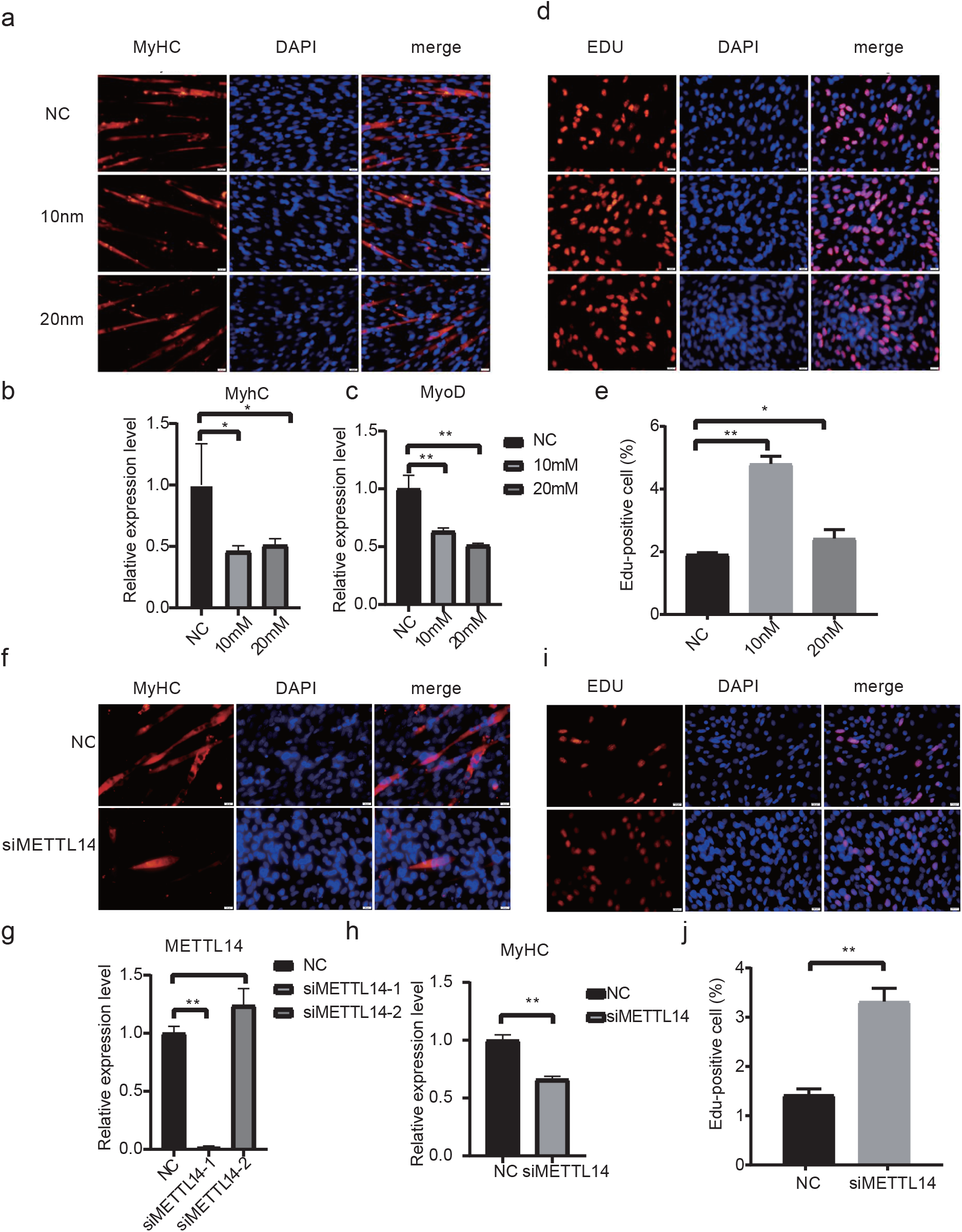
Suppression of m^6^A inhibited differentiation and promoted proliferation of myoblast cells. **(a, b)** MyHC immunofluorescence staining (20x) and RT-qPCR in C2C12 cells differentiated for 4 days showing that both 10nM and 20nM cycloleucine can reduce the protein and mRNA expression levels of MyHC as well as the mRNA expression level of MyoD **(c)**. **(d, e)** EdU staining showing that both 10nM and 20nM cycloleucine significantly increased EdU incorporation (20x). **(f, g, h)** MyHC immunofluorescence staining (40x) and RT-qPCR in C2C12 cells differentiated for 4 days showing that, in siRNA-medicated METTL14-knockdown cell, METTL14 expression was significantly decreased and METTL14-knockdown significantly reduced the protein and mRNA expression levels of MyHC. **(i, j)** EdU staining showing that METTL14-knockdown significantly increased EdU incorporation (40x). The data represent the means ± SD of three independent experiments. *P < 0.05, **P < 0.01.

### Generation of longitudinal transcriptomes and epi-transcriptomes of prenatal skeletal muscle

To further delineate the functional implications of m^6^A during skeletal muscle development, we then aimed to profile the transcriptome-wide m^6^A in prenatal skeletal muscle in pig. Due to the limited amount of prenatal skeletal muscle, a refined MeRIP (R-MeRIP) approach that can analyze only 5ug total RNA per sample was first tested in our study (Methods) [11], using a porcine PK15 cell line (Additional file 2:Table S1). Both the R-MeRIP and the conventional MeRIP sequencing were performed twice on the cell line and the m^6^A profiles generated were analyzed by the same analytical pipeline. As a result, a high correlation coefficient (R^2^~0.9) was achieved between the two independent tests for both the R-MeRIP and MeRIP methods, indicating for high reproducibility of both technologies (Additional file 1: Figure S1a). Furthermore, 66.22% of the m^6^A peaks called from two approaches were consistent (Additional file1: Figure S1b), as defined by a previously published algorithm (Methods).

Thereafter we used R-MeRIP to examine both transcriptomes and m^6^A epi-transcriptomes of prenatal skeletal muscle at six stages, including 33, 40, 50, 60, 70 and 95dpc, spanning two important waves of porcine myogenesis (Additional file 2: Table S2). Based on these data, m^6^A peaks were annotated for all the transcripts. As a result, we found mast majority (99.47%) of the peaks were in the coding genes (Additional file1: Figure S2a). Furthermore, a similar pattern of m^6^A distribution from 33pdc to 95dpc was observed, as these peaks of m^6^A were preferentially located at the coding sequence (CDS) and 3’ untranslated region (3’UTR) rather than the 5’UTR (Additional file1: Figure S2b, c), and the m^6^A peaks contained conservative sequence motifs of RRACH, in agreement with previous studies [5, 6] (Additional file1: Figure S2d). Despite of that, divergent numbers of m^6^A peaks and the methylated genes were revealed among different stages (Fig. 2a). Gene Ontology (GO) term enrichment analyses revealed that the m^6^A genes were commonly associated with RNA binding, nucleoplasm and macromolecule metabolic process, but relatively more divergent for functions of regulation of gene expression and biosynthesis processes (Fig. 2b). Furthermore, we also observed a gradual decrease of expressed genes across six stages (Fig. 2c) and clear classification of the gene expression program based on hierarchical clustering analysis (Fig. 2d). These results might imply for constant re-programming of epi-transcriptome during prenatal myogenesis that might be associated with the changed gene expression profiles in these samples.

**Fig. 2.**
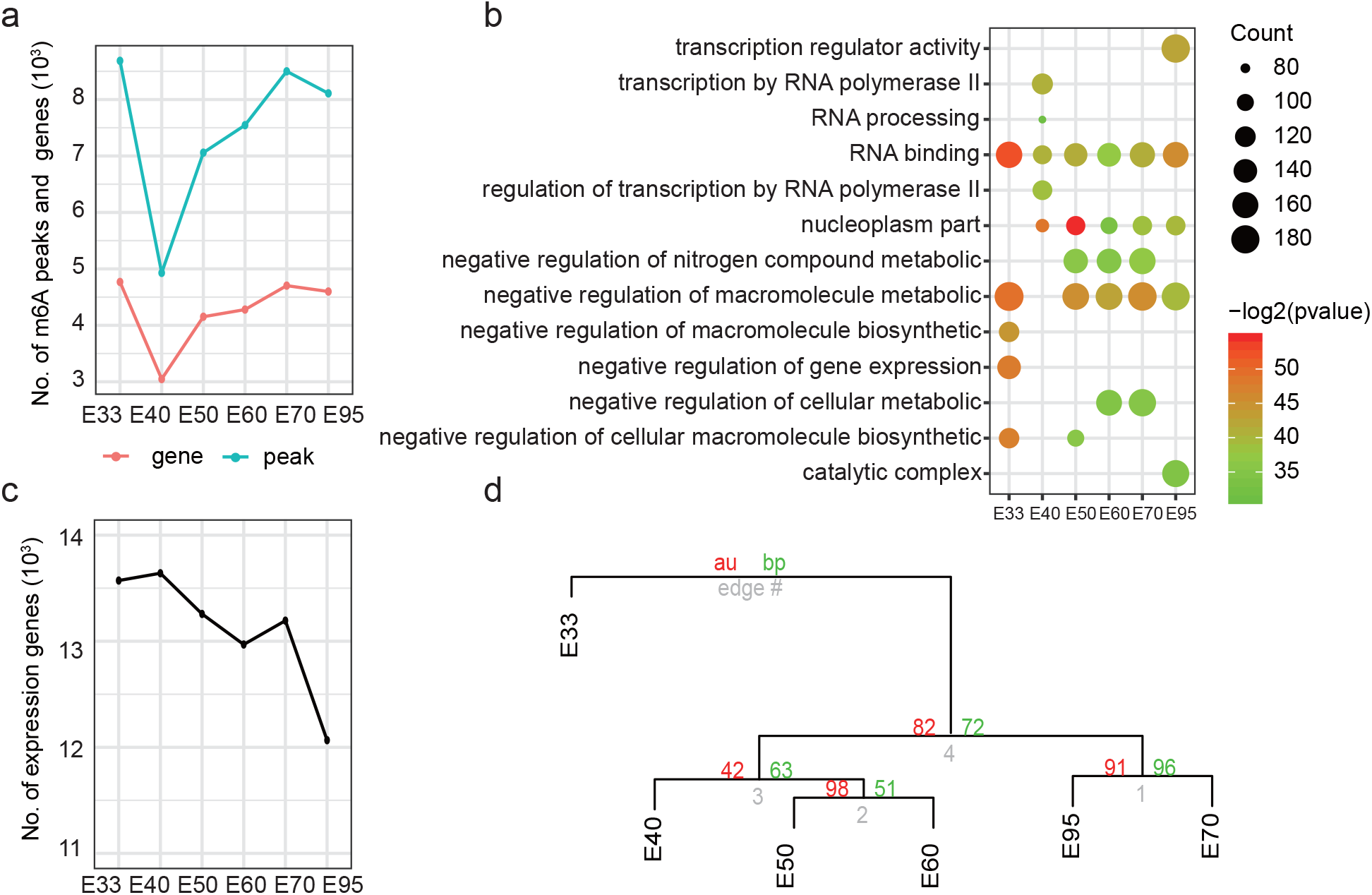
Characterizing the longitudinal transcriptome and epi-transcriptome of the six prenatal stages of skeletal muscle. **(a)** The number of genes containing m^6^A peaks and the number of m^6^A peaks in the six developmental stages. **(b)** Top 5 GO biological processes of m^6^A enriched genes in each of the six developmental stages. **(c)** The number of expressed genes (TPM >1 in each sample) in the six developmental stages. **(d)** Unsupervised hierarchical clustering of the six developmental stages based on transcriptomes. Abbreviations: E33: 33 day post coitus (dpc); E40: 40dpc; E50: 50dpc; E60: 60dpc; E70: 70dpc; E95: 95dpc.

### Dynamic transcriptome during porcine development of prenatal skeletal muscles

To investigate the genome-wide gene expression during prenatal skeletal muscle development, we further performed pair-wise comparisons between consecutive stages to reveal the differentially expressed genes (DEGs). As shown in Fig. 3a, interestingly, more DEGs were detected in comparison 33vs.40 and 70 vs.90 (3471 and 2312 DEGs, respectively) than in other comparisons (~500 DEGs) (Fig. 3a). These results of gene expression changes were in accordance with previous observations on the developmental events that landrace pigs showed the first onset of primary myofiber at 35dpc and an increased number of muscle fibers occurs between 77 and 91dpc in relation with the fusion of these myogenic cells [18]. To further identify key regulatory genes during muscle development, we used weighted gene co-expression network analysis (WGCNA) to detect all the DEGs identified across these stages of skeletal muscle development, with considering their correlation with body weight, body length and developmental stages [19]. We identified seven modules out of these DEGs, among which the magenta and darkgrey modules showed significant correlation with all phenotypes (p<0.05) (Fig. 3b). In the darkgrey module, the gene expression levels of DEGs were gradual increased from 33dpc to 95dpc, which were mainly associated with skeletal muscle development process (the top 5 significant biology processes). In contrast, the magenta module showed gradual decrease of expression levels, these genes were mainly involved in embryo and neuron system development (Fig. 3c). Among these genes, 5 and 3 genes expression were continuously up-regulated and down-regulated during the development (Fig. 3d), respectively. Notably, IGF2BP1 gene was continuously down-regulated, which is a key m^6^A reader that can regulate mRNA stability and translation [20] in the process of tissue/organ development [21]. These results again suggest that m^6^A may play a key role in prenatal myogenesis.

**Fig. 3.**
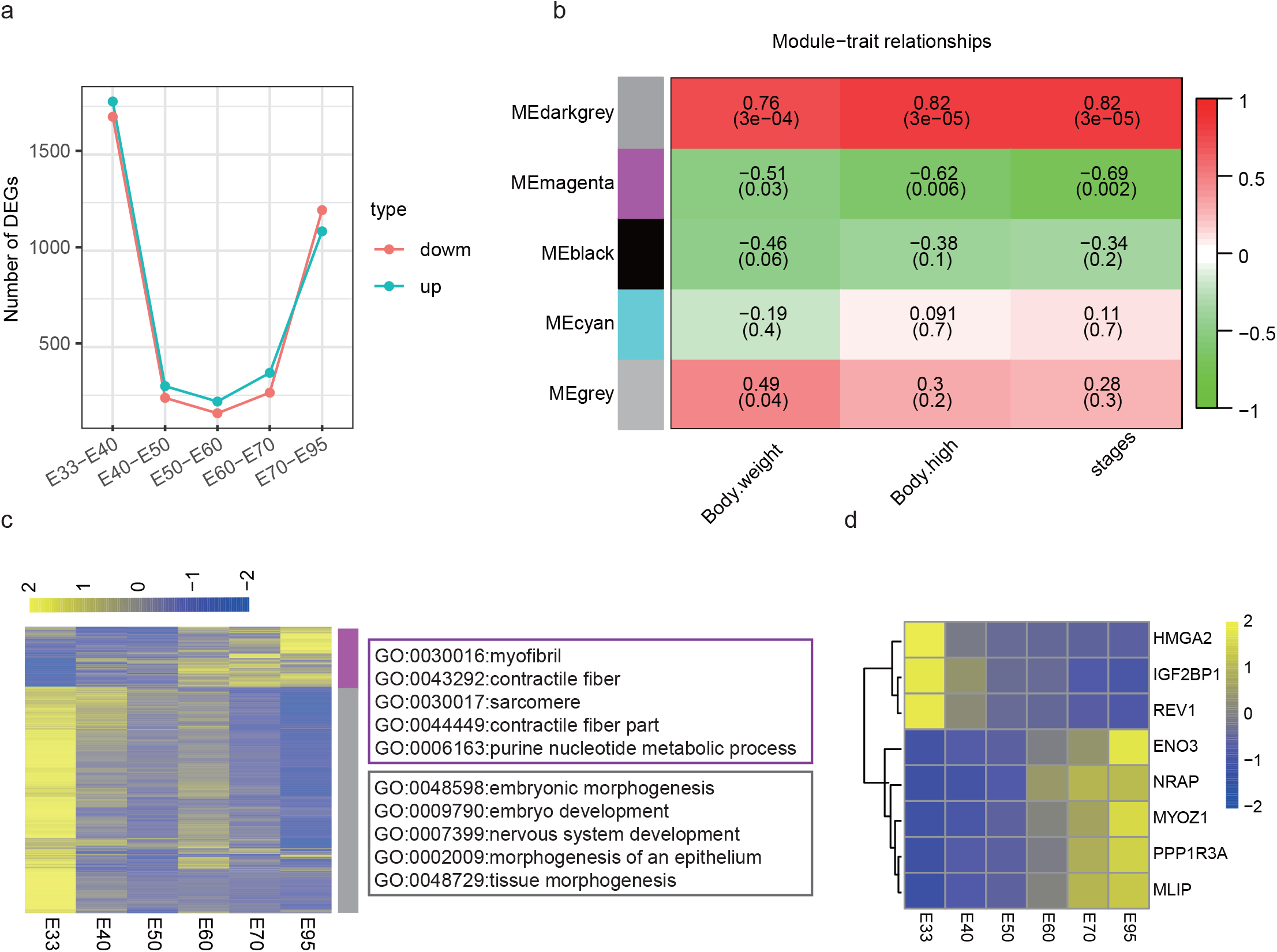
WGCNA analyses of expressed genes in the six prenatal stages of skeletal muscle. **(a)** The number of DEGs calculated from inter-stage comparisons. **(b)** Heatmap showing the correlation between module eigengenes and traits (each cell contained the corresponding correlation coefficients and P-values). **(c)** Heatmap showing the expression levels of genes in the darkgrey and magenta modules during the developmental process of prenatal skeletal muscle, top 5 GO biological process terms were indicated in the right side. **(d)** Heatmap showing the expression levels of genes that are either continuously up-regulated or down-regulated across the six developmental stages.

### m^6^A epi-transcriptome through developmental skeletal muscle

Next, we characterized the dynamic m^6^A modification during prenatal skeletal muscle development. We first combined the mRNA transcripts with m^6^A peaks. Majority (2466) of the methylated target were commonly methylated genes (CMG) throughout prenatal skeletal muscle, whereas only 414, 24, 95, 114, 244, and 312 specifically methylated genes (SMG) were identified at 33, 40, 50, 60, 70, and 95dpc, respectively (Fig. 4a). In comparison with SMGs, CMGs exhibited averagely more peak number and lower ratio of peaks located in 3UTRs (Additional file1: Figure S3a, b). A further unsupervised clustering of the CMGs based on the m^6^A levels led to two major clusters, for which GO analyses on CMGs indicated different profile of genes were enriched in transcriptional regulation of gene expression, such as regulation by RNA polymerase II, transcription regulator activity, and RNA processing (Fig.4b). In addition, a different enrichment of SMG genes was observed, more genes enriched in the pathways related with tissue genesis at early stage of embryo development (Additional file1: Figure S3c). To further explore the stepwise changes of m^6^A during prenatal skeletal muscle development, m^6^A peaks were subjected to pair-wise comparison between each two adjacent stages. In total, we identified 434 differentially methylated genes (DMGs), including 297 hyper-methylated genes and 137 hypo-methylated genes across six stages (Fig. 4c). Consistent with DEGs (Fig. 3a), the biggest dynamic m^6^A variation happened in the transition from 33dpc to 40dpc (Fig. 4c). Only a few genes were common between either two sets of inter-stage DMGs, indicating a highly dynamic changes of m^6^A during skeletal muscle development (Fig. 4d). GO analyses on each set of inter-stage DMGs displayed divergent enrichment of DMGs, also suggesting stage-specifically changes of dynamic m^6^A modifications during the muscle development (Fig. 4e).

**Fig. 4.**
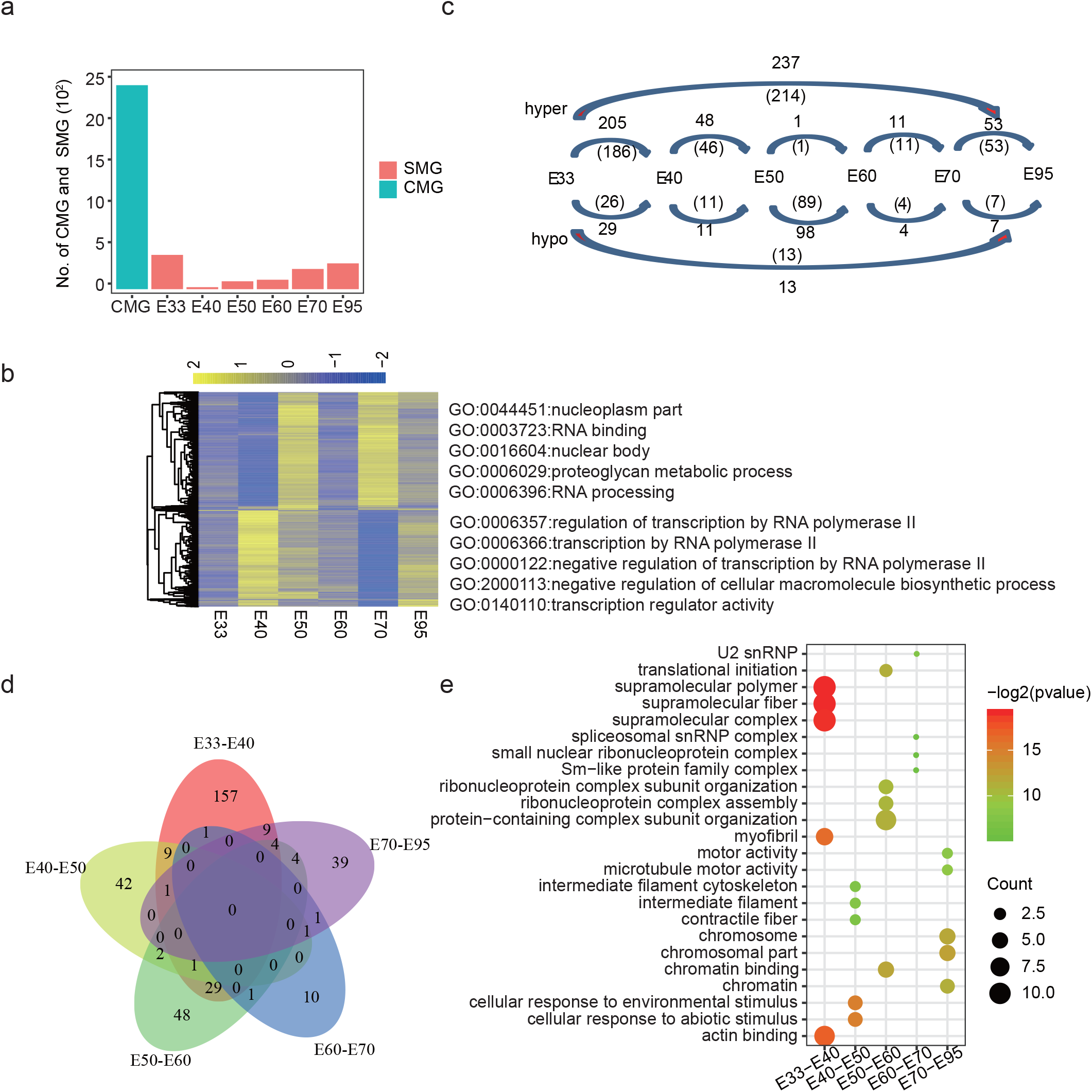
Charactering the dynamic m^6^A modification in the six prenatal stages of skeletal muscle. **(a)** Numbers of common (CMG) and temporal-specific (SMG) m^6^A genes in the six prenatal stages. **(b)** Heatmap showing the methylation level of CMGs that are unsupervised clustered during the developmental process of prenatal skeletal muscle, top 5 GO biological process terms were indicated in the right side. **(c)** Numbers of hyper-methylated or hypo-methylated DMGs from inter-stage comparisons across six prenatal stages. **(d)** Venn diagram showing the number of overlapped DMGs; **(e)** Top 5 GO biological process terms of each inter-stage DMG.

### m^6^A regulate gene expression of prenatal skeletal muscle

To further address the potential role of m^6^A in regulation of mRNA expression in prenatal myogenesis, we focused on the m^6^A variation in genes of darkgrey and magenta modules, which showed gradually increased or decreased expression over skeletal muscle development, respectively. Consistent with the changes of gene expression pattern, the ratio of m^6^A-modified genes in the total module gene-set (Fig. 5a) as well as the average peak number of the m^6^A-modified genes (Fig. 5b) presented similar tendency as of the mRNA levels in the darkgrey and magenta modules. We then cross-matched the DEGs from these two modules with the 434 identified DMGs, resulting 97 and 30 overlapped genes from the magenta and darkgrey module, respectively (Fig. 5c). These 127 genes were further filtered for those genes contained both significantly differential m^6^A and transcripts at the same stage and classified them into two categories based on whether the inter-stage fold changes of m^6^A or gene expression displayed the same (27 genes) or opposite (42 genes) tendency. Furthermore, correlation analysis indicated the correlation between the fold changes of m^6^A and gene expression for both categories was significant, either positively or negatively (Fig. 5d; Additional file 1: Figure S4a.). These genes, were largely enriched in GO terms for muscle development (Fig. 5e, Additional file 1: Figure S4b). Especially, we found that the positively correlated genes (Additional file 2: Table S3) were mainly enriched in the darkgrey module from two inter-stages (33 to 40dpc, 70 to 95dpc), whereas most of the negatively correlated genes (Additional file 2: Table S4) were from 33 to 40dpc (Fig 5e, Additional file 1: Figure S4b). Together, these results confirmed that m^6^A was involved in the regulation of gene expression during prenatal skeletal muscle development. As one example gene, MYH1 (Myosin-1) as an isoform consist of Myosin heavy chain (MyHC) [22], significantly elevated its m^6^A and expression levels from 40dpc to 50dpc and 70dpc to 95dpc (Fig. 5e).

**Fig. 5.**
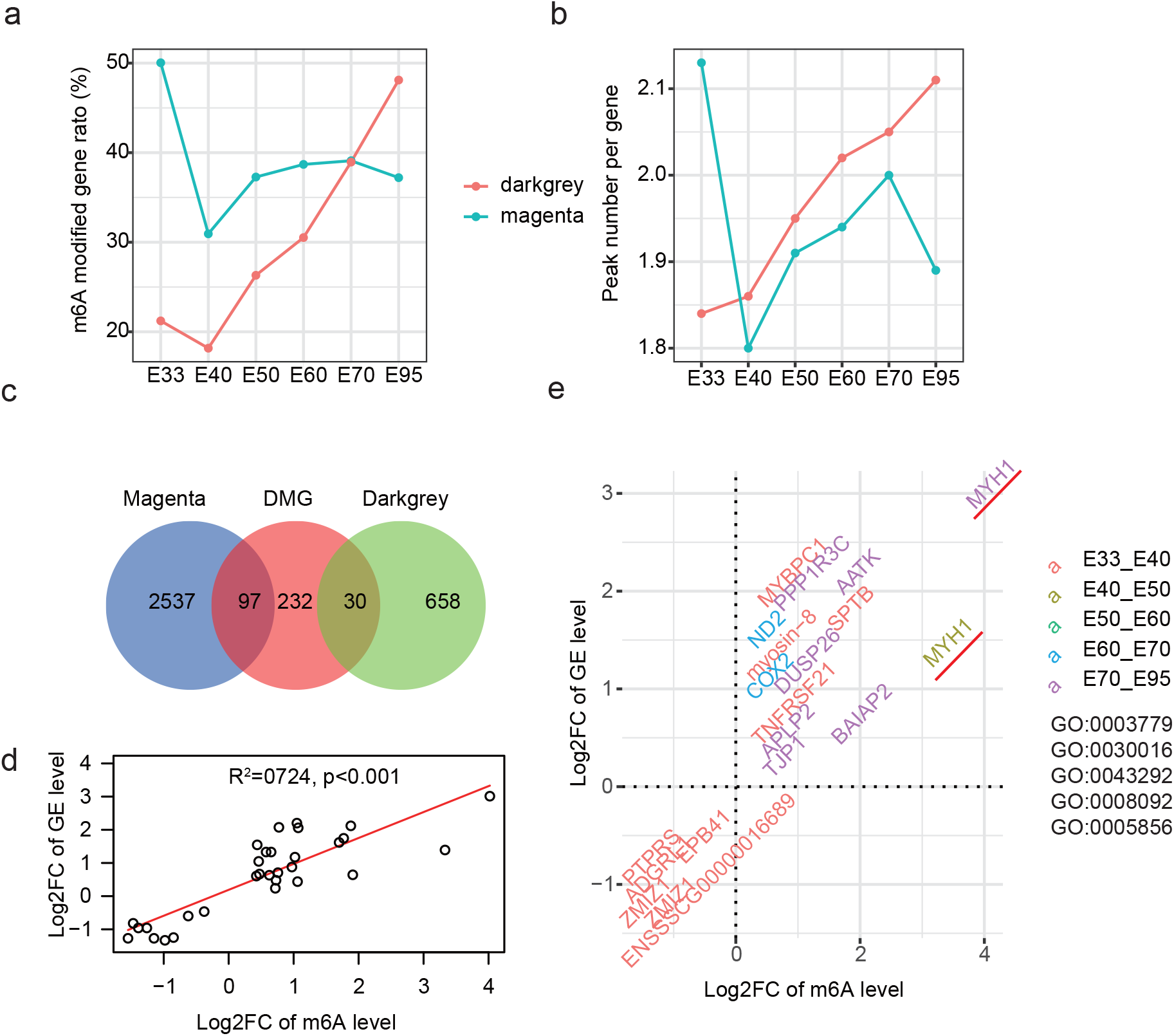
m^6^A characteristics of genes in the two modules that showed significant correlation between gene expression levels and phenotypes. m^6^A modified gene ratio **(a)** and average peak number of all m^6^A genes **(b)** for the darkgrey and magenta modules gene in each developmental stage. **(c)** Venn diagram showing the overlapping of two significant correlation module genes and DMGs. **(d)** Correlation between the fold changes (FC) of m^6^A and gene expression (GE) for same tendency gene; part genes and top 5 GO biological process terms was displayed in **(e)** (GO:0003779: actin binding; GO:0030016: myofibril; GO:0043292: contractile fiber; GO:0008092: cytoskeletal protein binding; GO:0005856: cytoskeleton).

### IGF2BP1-dependent m^6^A regulates myoblast proliferation and differentiation

Previous studies indicated that m^6^A can regulate RNA stability (positive correlation) or promote m^6^A degradation (negative correlation) with help of reader proteins IGF2BP1 and YTHDF2, respectively. The significantly dynamic changes of the gene expression of IGF2BP1 across the six stages also indicated its role in myogenesis (Fig. 3d). To further confirm whether the reader protein IGF2BP1 participates in regulation of myogenesis, therefore, we knocked down it in C2C12 myoblast (Fig. 6a). We found that the knockdown of IGF2BP1 gene significantly decreased the expression of MyHC at protein and mRNA level (Fig. 6b, c), whereas obviously increased the EdU incorporation (Fig. 6d, e). These results indicated that knockdown of IGF2BP1 suppressed differentiation and promoted cell proliferation of C2C12 myoblast, consistent with the results of chemical suppression m^6^A and knockdown METTL14 treatments.

**Figure 6.**
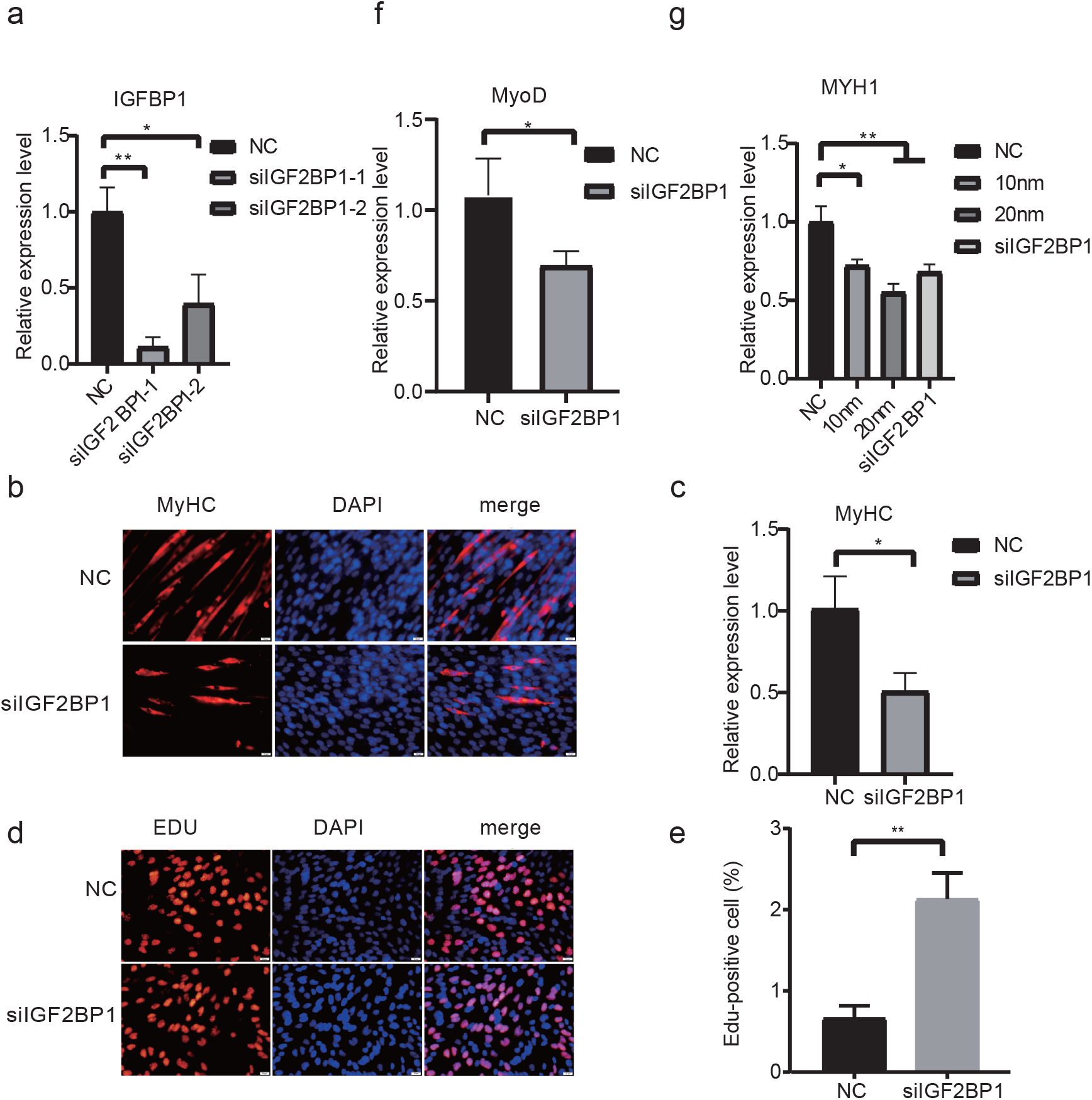
IGF2BP1-knockdown inhibits myoblast differentiation. **(a)** IGF2BP1 expression was significantly reduced in siRNA-mediated IGF2BP1-knockdown C2C12 cell. **(b, c)** Photographs of MyHC immunofluorescence staining (20x) and RT-qPCR in C2C12 cells differentiated for 4 day showing that IGF2BP1-knockdown inhibited protein and RNA level of MyHC. **(d, e)** Photographs of EdU staining showing that IGF2BP1-knockdown significantly increased C2C12 EDU incorporation (20x). (**f, g**) The mRNA expression level of MyoD and MYH1 was detected by RT-qPCR. The data represent the means ± SD of three independent experiments. *P < 0.05, **P < 0.01.

Furthermore, considering the changed expression of MyoD and MyHC in response to suppression of m^6^A and knockdown of METTL14 treatment (Figure 1), we also examined the expression of MyoD in C2C12 cell with knocked-down IGF2BP1. Quantitative results showed that their expression was significantly reduced, which raised the possibility that m^6^A regulate the expression of MyoD through an m^6^A-IGF2BP1-dependent pathway (Fig. 6f). In addition, we examined the RNA expression of MYH1 that are positively regulated by m^6^A. MYH1 expression was significantly decreased in C2C12 cells upon knockdown of IGF2BP1 (Fig. 6g), suggesting that IGF2BP1 might play an important role in the regulation of MYH1.

## Discussion

Skeletal muscle development is known to be regulated in a complex spatiotemporal way. Previously identified changes in transcriptional expression and epigenetic modification throughout skeletal muscle development have greatly improved our understanding of the mechanism related to myogenesis [3, 23]. In present study, we hypothesized that m^6^A might also participate in skeletal muscle development, given the role of m^6^A participating in regulation of tissue development, including cerebellum, neural and fat [24–26]. By using methodologies of either knockdown of key m^6^A methylase or binding proteins in C2C12 and high-throughput sequencing of m^6^A profiles across developmental skeletal muscle at six stages, we demonstrated that m^6^A modification plays an important role in myogenesis, especially affecting myoblast proliferation and differentiation (Fig. 1, Fig. 6). Across the developmental skeletal muscle, a highly dynamic m^6^A epi-transcriptome as well as transcriptome were revealed, in line with other studies on the regulatory role of m^6^A in tissue development, e.g., postnatal liver development in pig [27] and mouse cerebellum development [25]. Especially, we observed that the most divergent stage interval during the two important waves in prenatal fiber formation is from 33 to 40dpc, which is in the period of primary fiber formation. As previously confirmed, the gene expression changes was also more intense in the period of primary fiber formation than in the period of secondary fiber formation [28]. These results imply m^6^A also participates in the developmental regulation of prenatal skeletal muscle.

Previous studies suggested that m^6^A modification can maintain RNA stability with help of a m^6^A reader protein IGF2BP1 [29] [20]. The myosin immunofluorescence and EDU staining assay indicated that IGF2BP1 also participates in the regulation of myoblast proliferation and differentiation (Fig. 6). During prenatal skeletal muscle development, we found the transcription of IGF2BP1 continuously decreased. Thereby, we revealed a set of genes with significantly positive correlation between the inter-stage fold changes of m^6^A and gene expression. Among these genes, MYH1 codes for an isoform of Myosin heavy chain (MyHC), and twice appear in positive correlation gene list (Fig. 5d). The level of MYH1 was significantly decreased in C2C12 myoblast under the cycloleucine treatment and IGF2BP1-knocked down (Fig. 6g). Whether IGF2BP1 directly binds to these key genes in skeletal muscle development requires further examination. Over the past years, several studies have unraveled important mechanisms by MyoD, control the specification and the differentiation of the muscle lineage [30, 31]. The Myosin heavy chains (MyHC) are terminally differentiated muscle cells markers [16]. Both chemical suppression of m^6^A and knockdown METTL14 decrease the expression of MyoD and MyHC (Fig. 1) in C2C12, which was consistent with the regulation of IGF2BP1. Therefore, we evaluated the involvement of IGF2BP1, which maintains RNA stability. Our result showed that knockdown IGF2BP1 also significantly reduce the MyHC and MyoD at RNA level (Fig. 6c, f). MyoD was m^6^A-modified target in C2C12 [32]. These results suggest that m^6^A regulate MyoD at RNA level may be through an m^6^A-IGF2BP1-dependent pathway, which may be the main reason for m^6^A regulate myoblast differentiation.

m^6^A modification modulates all stages in the life cycle of RNA, such as RNA processing, nuclear export, and translation modulation [33, 34]. First, YTHDF2 trigger degradation through conveying m^6^A mRNA to specialized mRNA decay machineries (P bodies etc.) [35]. Second, YTHDF1 protein promote synthesis by interacting with translation machinery [36]. Although translation and degradation are two opposite fates of mRNA, two reader protein share percent of fifty common target mRNAs [36]. m^6^A possess a multi-dimensional mechanism of mRNA methylation in modulating gene expression. IGF2BPs promote the stability and storage of their target mRNAs in an m^6^A-dependent manner under normal and stress conditions [36]. As YTHDF2, IGF2BPs is highly likely located in processing bodies (P-bodies) [20]. Our result show that only one gene was overlapped between positive and negative correlation gene (Additional file1: Figure S4c). IGF2BPs and YTHDF2 have a distinct pattern of binding sites [20]. Published binding gene of IGF2BP1 and YTHDF2 have nearly 20% common target (Additional file1: Figure S5a). IGF2BP1 and YTHDF2 may regulate their own subsets of mRNA targets, independently. Furthermore, IGF2BPs also enhance mRNA translation [20]. More than percent of fifty YTHDF1 targets is also IGFBPs targets (Additional file1: Figure S5b). A temporal order may exist for the regulation of two reader proteins common target gene translation. Further work should attempt to uncover the relationship between IGF2BP1 and YTHDF2/YTHDF1on m^6^A modification gene.

## Conclusions

Here, we describe m^6^A RNA methylation landscape during prenatal development of the porcine skeletal muscle and verify the function of m^6^A methylation in C2C12 myoblast. Our study reveals the critical role of m^6^A methylation in prenatal skeletal muscle development. Reader protein IGF2BP1 participates in the regulation of myoblast proliferation and differentiation by affecting the expression of MyoD, MyHC and MYH1. These findings highlight the function of m^6^A methylation in prenatal skeletal muscle development.

## Material and Methods

### Cell culture

C2C12 cells were grown in incubators at 37°C and 5% CO_2_, and proliferating cells were cultured in Dulbecco’s Modified Eagle’s Medium (DMEM) supplemented with 10% fetal bovine serum (FBS; Gibco, Grand Island, NY, USA). For myogenic differentiation, cells were transferred to DMEM containing 2% horse serum (HS; Gibco). All cells were grown to 80-90% confluence before differentiation was induced.

### Animal and sample collection

All animal procedures were performed according to protocols approved by Hubei Province, PR China for Biological Studies Animal Care and Use Committee. Landrace was mated with the boar breed. The sows were then sacrificed at a commercial slaughterhouse at six stages (33, 40, 50, 60, 70, 95dpc). The uteri containing the fetuses were collected immediately, and the *longissimus* muscle tissues were rapidly and manually dissected from each fetus. At each time point, three randomly chosen embryos were used for the following research. In total, 18 samples were snap-frozen in liquid nitrogen and stored at –80°C until further use.

### siRNA synthesis and cell transfection

The siRNA sequences of METTL14, IGF2BP1 and YTHDF2 were synthesized as follows: METTL14 siRNA (sense): CAGUACCUUUCUUAAGGGATT; METTL14 siRNA (antisense): UCCCUUAAGAAAGGUACUGTT. IGF2BP1 siRNA (sence): GAGCACAAGAUCUCCUACATT; IGF2BP1 siRNA (antisence): UGUAGGAGAUCUUGUGCUCTT. For cell transfection, C2C12 cells were transfected approximately 10μl of siRNA oligo using Lipofectamine 2000 (Invitrogen) in each well of a 6-well plate.

### EdU staining and Immunofluorescence staining

Cell Immunofluorescence staining was performed according to the previously published method [37]. Immunofluorescence staining antibodies included MyHC (sc-376157; 1:200; Santa Cruz Biotechnology, USA), and a secondary antibody (antimouse CY3; Beyotime Biotechnology, China). DAPI was used to visualize the cell nuclei with a fluorescence microscope (DP80; OLYMPUS, Tokyo, Japan).

### qPCR

Total RNA was extracted using the TRIzol reagent (Invitrogen) and then reverse transcribed using HiScript III RT SuperMix for qPCR (Vazyme R323-01). qPCR analysis was performed using the Applied Biosystems StepOnePlus RealTime PCR system. All primers used in the study are presented in Additional file 2: Table S5. The relative RNA expression levels were calculated using the Ct (2–ΔΔCt) method [38].

### MeRIP sequencing

Total RNA was immediately isolated using Trlzol Reagent according to the manufacturer’s instructions (ambion). To avoid DNA contaminations all samples were treated with Dnase (NEB). m^6^A profiling for each individual was performed as previously described, respectively [5]. For isolation of mRNA, total RNA was subjected to two rounds of purification using oligo (dT)-couple magnetic beads according to the manufacturer’s instruction (Invitrogen). mRNA sample was fragmented into ~100nt fragments by 1min incubation at 90°C in fragmentation buffer (Invitrogen). About 50ng fragments mRNA samples were used to construct the input library. Other mRNA fragments were incubated for 4h at 4°C with 3ug anti-m^6^A polyclonal antibody (Synaptic systems, 202003), which has combined with protein-A beads (Invitrogen) at room temperature for 1 h in IPP buffer (150nm NaCl, 0.1% NP-40,10nm Tris-HCl). Bound mRNA was eluted from the beads with 0.5mg/ml N6-methyladenosine in IPP buffer. Eluted mRNA was precipitated by ethanol-NaAc solution and glycogen (Life Technologies) overnight at −80°C. mRNA was resuspended in H_2_O and used for library generation. IP mRNA and input mRNA respectively conducted an RNA-seq library using a Vazyme mRNA-seq kit (Vazyme NR601). Sequencing was carried out on Illumina HiSeq X ten sequencing.

### R-MeRIP sequencing

Total RNA was fragmented to 100-150 nucleotides by 15min incubation at 70°C in fragmentation buffer (Invitrogen). About 10ng fragments RNA was reserved to conduct the input library. Other RNA was immunoprecipitated with m^6^A antibody as MeRIP-seq to get m^6^A RNA. Once successfully immunoprecipitated methylated RNA was confirmed, input and IP library were constructed using SMARTer® Stranded Total RNA-seq Kit (Takara, 635007), respectively. This kit removes ribosomal cDNA using probes specific to mammalian rRNA post reverse transcription and amplification. Sequencing was also carried out on Illumina HiSeq X ten sequencing.

### RNA-seq data analysis

For each sample, pair-end reads were used for bioinformatics analysis. Quality control of raw data was done using FastQC software (version 0.11.3). Sequencing data were trimmed by Trimmomatic 0.36 [39] to remove the adapter and low-quality data. These high-quality reads were mapped against the ensemble pig genome (Sscrofa11.1) using hisat2 software (version 2.0.4) [40]. The TPM (Transcripts per million reads) values of RNAs in each sample were calculated using Salmon [41]. Gene TPM >1 in each sample was identified as expression gene. The differentially expressed gene (DEG) was identified using DESeq2 [42]. WGCNA (v1.66) was used to construct the unsigned co-expression networks based on gene expression [43]. All software applications were run with default parameters.

### m^6^A peak calling and motif analysis

RNA m^6^A-modified regions (m^6^A peaks) were identified using exomePeak software (version 2.14.0) [44, 45]. To improve m^6^A peak identification, consistent peaks in three biological replicates identified with exomePeak was regarded as highly enriched m^6^A peak for further analysis (P-value <0.05). The consensus sequence motif enriched in m^6^A peaks were identified by MEME [46].

### Characterization of m^6^A peak distribution patterns

Distribution of m^6^A peaks along mRNAs were obtained as previously [6], a reference porcine transcriptome was built using the longest transcript of each gene. Each of the 5’UTR, CDS, 3’UTR regions were splint into 100 bins with equal length. The percentage of m^6^A peaks in each bin was calculated to represent the occupancy of m^6^A peak along the whole transcripts.

### GO analysis

The clusterProfiler R package was used for GO analysis by applying the default parameter [47]. Plot was generated based on the enriched GO terms using ggplot2 in R package. Color intensity indicates that the value of –log2 (p-value), while the size of the circle indicates the gene counts.

## Data availability

The high-throughput m^6^A sequencing data have been deposited in the GEO database under the accession code GSE141943, m^6^A peaks in the six developmental stage are included in processed data.

## Authors’ contributions

Conception and design: FG, ZT; acquisition of the data: XZ; analysis and interpretation of the data: FG, XZ, JH, YY; cell proliferation, differentiation, transfection experiments: YY, YC; writing of the paper: FG, XZ.

## Competing interests

The authors declare that they have no competing interests.

## Acknowledgments

FG was supported by the Agricultural Science and Technology Innovation Program and The Elite Young Scientists Program of CAAS. ZT was supported by National Natural Science Foundation of China (31830090), the National Key Project (2016ZX08009-003-006), the Shenzhen Dapeng New District Special Fund for Industry Development (KY20180114) and the Agricultural Science and Technology Innovation Program (ASTIP-AGIS5).

