## Additional file 1 for "Longitudinal epi-transcriptome profiling reveals the crucial role of m^6^A in prenatal skeletal muscle development of pigs"

**Figure S1. Evaluation of R-MeRIP technology in comparison with MeRIP**

**Figure S2. Characterizing m6A epi-transcriptomes of porcine prenatal myogenesis process**

**Figure S3. Characterizing the common and specific m6A peaks across the six developmental stages**

**Figure S4. Characterizing genes with negative correlation between the fold change of m6A and gene expression**

**Figure S5. Target genes of three binding proteins of m6A**

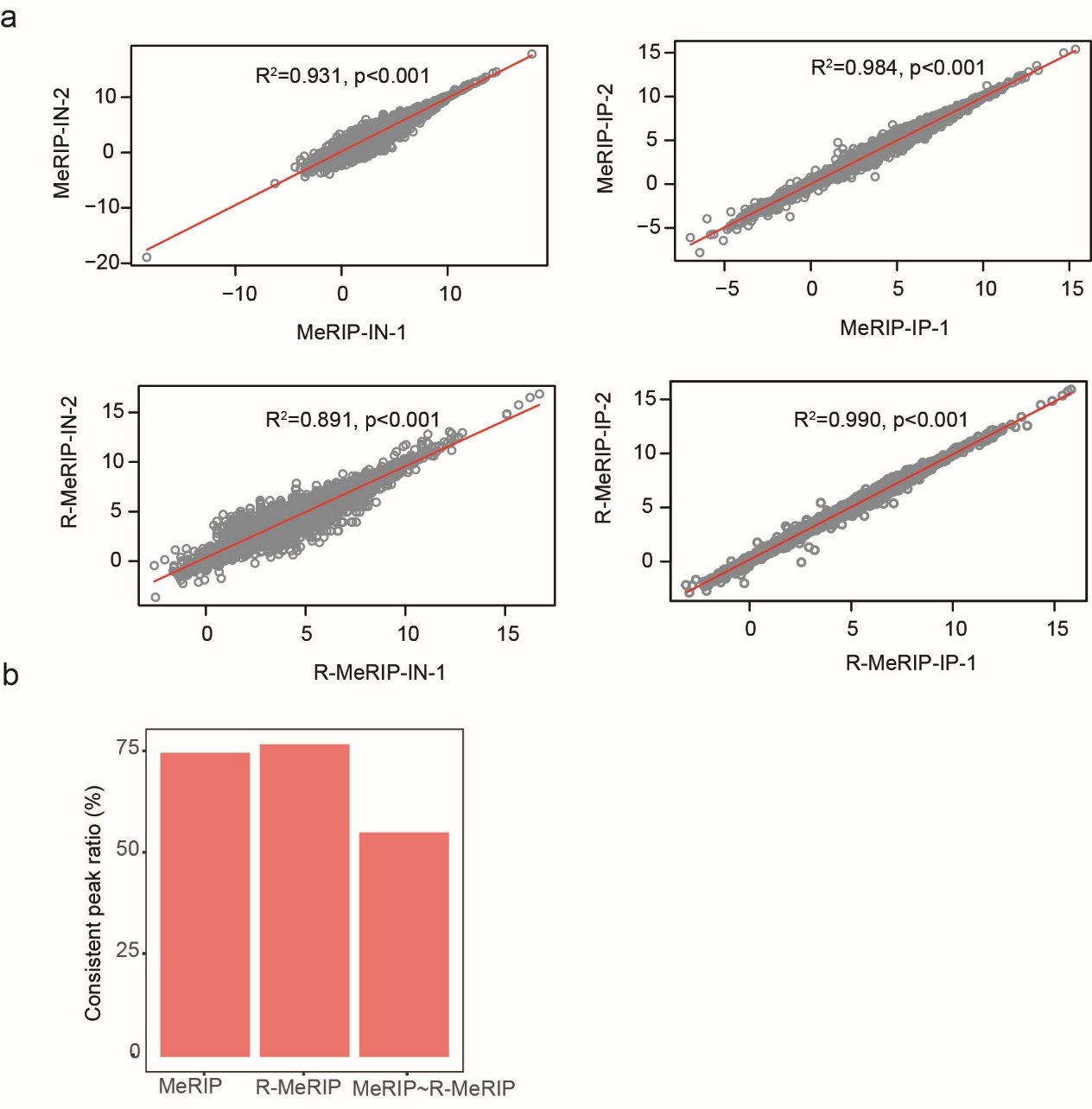

**Additional file 1： Figure S1 Evaluation of R-MeRIP technology in comparison with MeRIP**

**(a)** Correlation between two replicates of both the Input (IN, left) and IP (IP, right) libraries for MeRIP-seq (top) and R-MeRIP-seq (bottom) data**. (b)** Ratio of consistent peaks between two replicates of MeRIP or R-MeRIP and between MeRIP and R-MeRIP. The evaluation of consistent peak was done by exomePeak [1, 2]

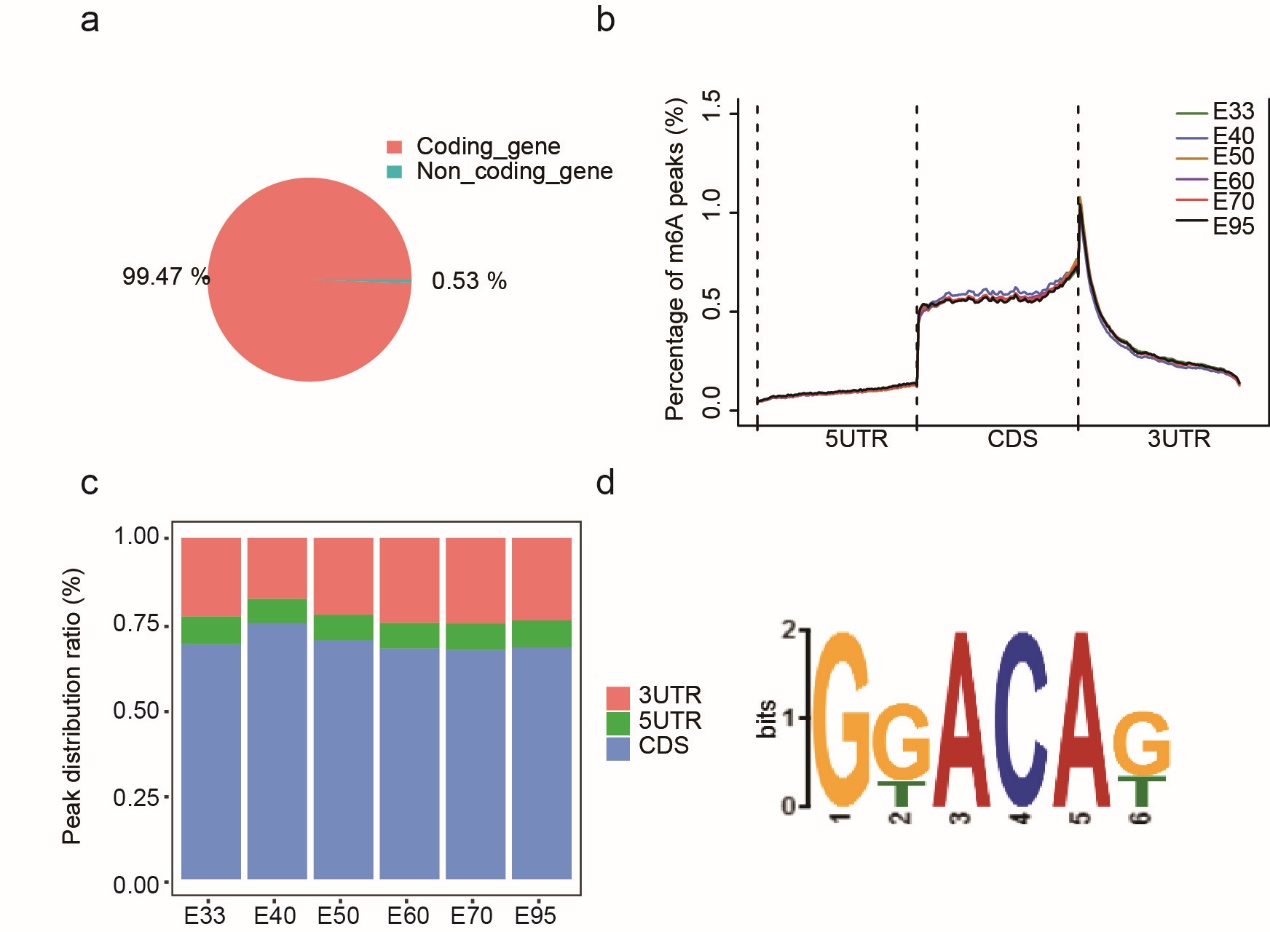

**Additional file 1： Figure S2 Characterizing m6A epi-transcriptomes of porcine prenatal myogenesis process**

**(a)** Distribution of m6A peaks on coding and non-coding gene; **(b)** Distribution of each stage m6A across the whole transcript; **(c)** The percentage of m6A peaks located in each categories of the transcript, i.e. 5UTR, CDS, and 3UTR; **(d)** Consensus sequence of the m6A peaks identified for porcine skeletal muscle.

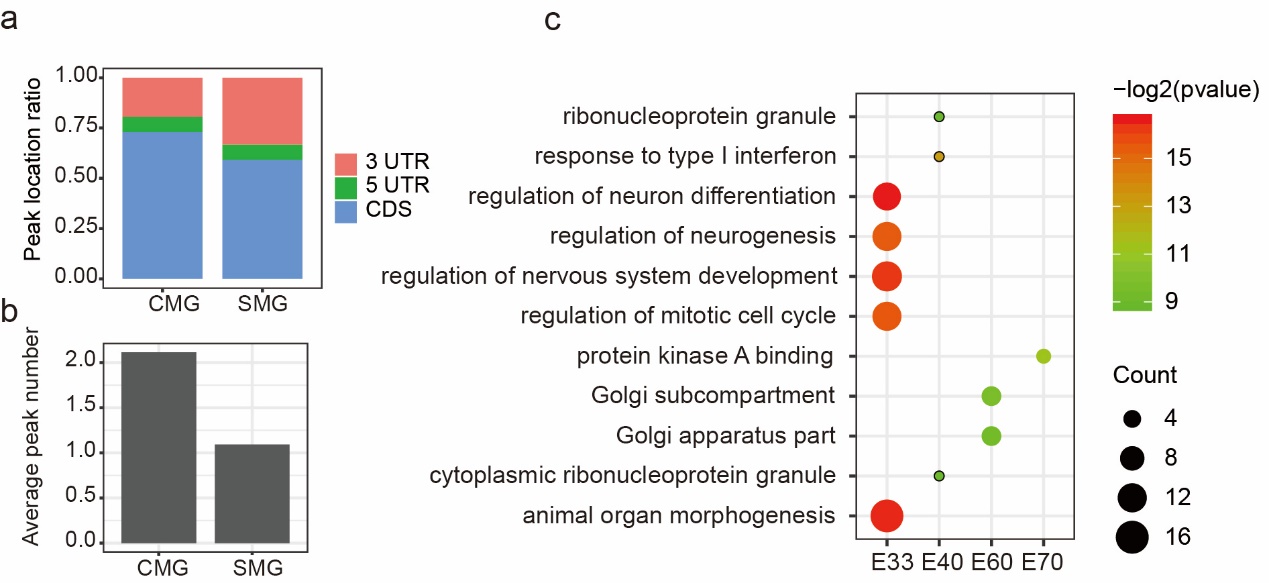

**Additional file 1： Figure S3 Characterizing the common and specific m6A peaks across the six developmental stages**

**(a)** The percentage of commonly methylated gene (CMG) and specifically methylated gene (SMG) identified across the six developmental stages; **(b)** Average peak number per gene for CMG and SMG; **(c)** GO terms of the enriched biological processes for SMGs in E33, E40, E60 and E70. No enrichment was revealed for E50 and E95.

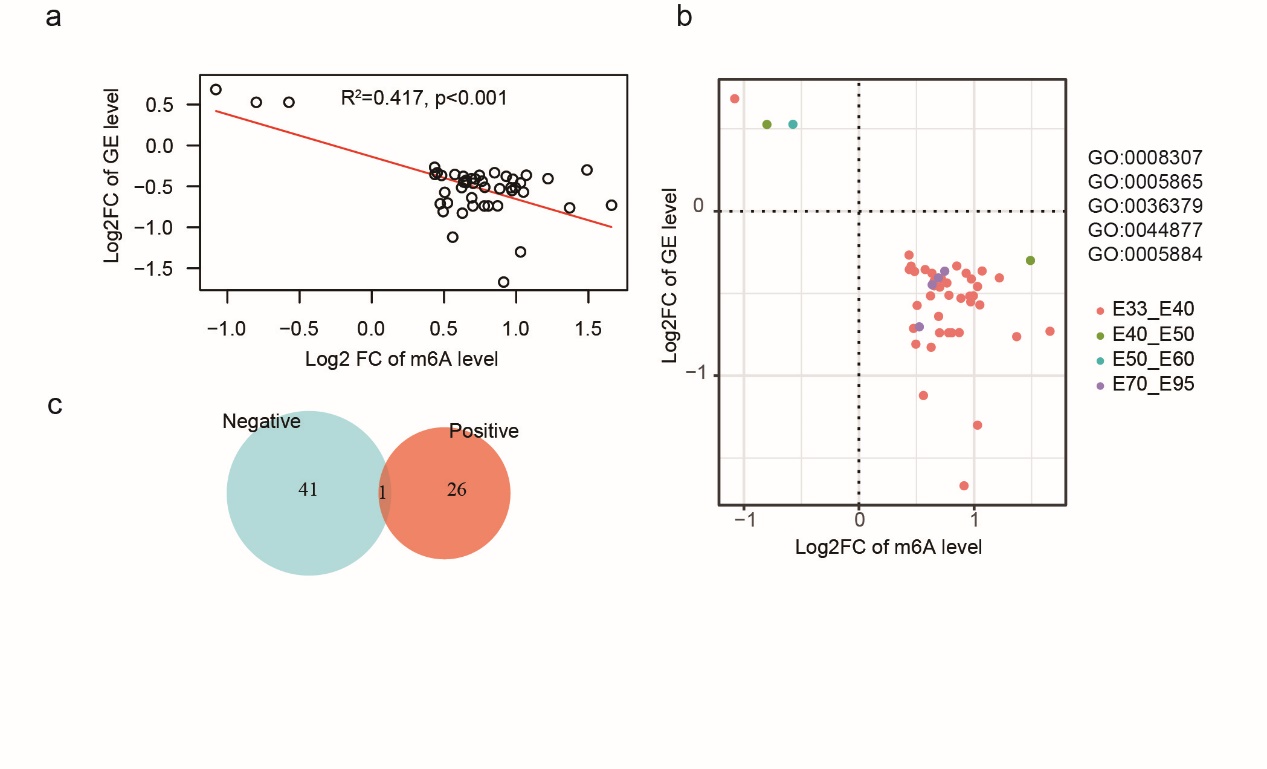

**Additional file 1： Figure S4 Characterizing genes with negative correlation between the fold change of m6A and gene expression**

**(a)** Negatively correlation between the fold changes of m6A and gene expression for opposite tendency genes; stage and top 5 GO biological process terms were displayed in **(b)** (GO:0008307: structural constituent of muscle; GO:0005865: striated muscle thin filament; GO:0036379: myofilament; GO:0044877: protein-containing complex binding; GO:0005884: actin filament); **(c)** Venn diagram showing the overlapping between positive and negative correlation gene.

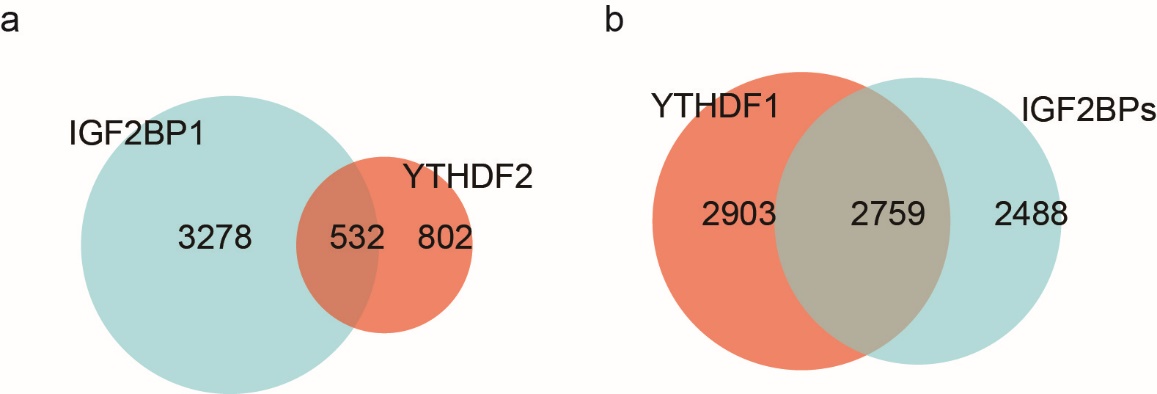

**Figure S5 Target genes of three binding proteins of m6A**

**(a)** Venn diagram showing the overlapping of target genes of IGF2BP1 and YTHDF2; **(b)** Venn diagram showing the overlapping of target genes of IGF2BPs and YTHDF1.
